# Goalkeeper game: a new assessment instrument in neurology showed higher predictive power than moca for gait performance in people with parkinson’s disease

**DOI:** 10.1101/400457

**Authors:** Rafael B. Stern, Matheus d’Alencar, Yanina L. Uscapi, Marco D. Gubitoso, Antonio C. Roque, André F. Helene, Maria Elisa P. Piemonte

## Abstract

**Objective:** To investigate the use of the Goalkeeper Game (GG) to assess gait automaticity decline under dual task conditions in people with Parkinson’s disease (PPD) and compare its predictive power with the one of the MoCA test.

**Materials and Methods:** 74 PPD (H&Y stages: 23 in stage 1; 31 in stage 2; 20 in stage 3), without dementia (MoCA cut-off 23), tested in ON period with dopaminergic medication were submitted to single individual cognitive/motor evaluation sessions. The tests applied were: MoCA, GG, dynamic gait index (DGI) task and timed up and go test (TUG) under single and dual-task (DT) conditions. GG test resulted in 9 measures extracted via a statistical model. The predictive power of the GG measures and the MoCA score with respect to gait performance, as assessed by DGI and DT-TUG, were compared.

**Results:** The predictive models based on GG measures and MoCA score obtained, respectively, sensitivities of 65% and 56% for DGI scores and 59% and 57% for DT-TUG cost at a 50% specificity. GG application proved to be feasible and aroused more motivation in PPDs than MoCa.

**Conclusion:** GG, a friendly and ludic game, was able to reach a good power of gait performance prediction in people at initial and intermediate stages of PD evolution.

## 1 Introduction

Parkinson’s disease (PD) daily living independence decrease and, consequently, the reduction in Health-Related Quality of Life (HRQoL) [40], are directly associated to deficiency in automaticity. One of the most devastating deficiencies involves gait impairments, product of inability of the nervous system to successfully coordinate movement, driving patients to a gradual increase in cortical demand to execute basic motor operations via attentional processes [6, 64, 13].

Although gait description typically involves an implicit ability that can be automatically performed, during tasks involving complex motor and perceptual integration, gait performance can become dependent on both automatic (implicit) and attentional (explicit) guided processes [65, 67, 2]. As in people with PD (PPD), even the gait parameters controlled by automatic processes in healthy individuals become dependent on attentional control [64, 12, 37], simple tasks become complex actions and complex ones, as walking over obstacles [53, 19, 1], avoiding obstacles [43], or managing unexpected targets and obstacles [8], can be impracticable. Moreover, assuming that each individual has a certain maximum attentional reserve capacity, when more attentional resources are needed to compensate for the deficits in automaticity, less of these resources remain to be used for other simultaneous tasks. The impact of this tradeoff can, sometimes, hide actual gait deficiency but is always noticeable in the impact of a concurrent cognitive task in PPD [50].

In fact, dual task (DT) gait paradigm, e.g., walking while performing a second task, is commonly applied to investigate gait automaticity in PD. DT gait paradigm allows the assessment of effects of attention division, permitting the detection of gait deficits that under single task (ST) (walking with no other assigned task) condition could be undetectable [7]. Gait evaluation under DT has been stated as a reliable measure for clinical studies in PD [55]. The change in gait performance under DT conditions relative to the ST condition is called dual-task cost (DTC) [13]. DTC has been largely demonstrated to be a reliable measure of impaired DT gait performance in PD [65, 66, 44, 47, 3, 25, 59].

The association between cognitive and motor disability performances provides a basis for understanding the complex role of cognition in Parkinsonian gait. The impaired ability of PPD to adapt the stepping and walking behavior towards targets and obstacles has significant correlations with the Montreal Cognitive Assessment (MoCA) score, which may reflect less effective behavioral responses due to attentional control deficits and/or impaired cognitive function [45]. In clinical routine, MoCA test is widely used to evaluate the cognitive status in PPD [11, 22, 29, 42, 51], being able to detect alterations even in early stages of the disease [33]. The MoCA test is a useful screening tool for global cognitive and executive functions in PD [27]. However, DT costs combined with clinical confounders were able to explain only about 30% of the MoCA score variance [20].

New tools may provide alternative and sensitive methods of early and non-invasive automaticity decline screening in PD. Identification of early automaticity impairment could provide a critical opportunity for early intervention before gait changes with major impact on independence in daily living activity, fall risk, and HRQoL. The Goalkeeper Game (GG) was introduced in de Castro [15] as a tool to investigate the conjecture that the brain does statistical model selection. GG is a videogame with internet, desktop and mobile device versions (http://game.numec.prp.usp.br) in which the player, taking the role of a goalkeeper in a soccer penalty shootout, guesses the position in the goal that the ball will hit (left side, right side or center). The game consists in a sequence of penalty kicks in which the ball positions can be generated either deterministically or randomly according to a strategy described by a tree and unknown to the player. The strategy is fixed for each phase and as the player (the goalkeeper) succeeds in guessing enough hits, which depends on the strategy tree, the phase terminates and a new one starts with a more complex tree. As the game evolves, the expectation is that for large numbers of trials in each phase the player is able to make sense of the strategy and obtain a high-scoring performance. Currently, the GG is being used by the Research, Innovation and Dissemination Center for Neuromathematics (http://neuromat.numec.prp.usp.br/) as an assessment tool in its basic and applied neuroscience researches.

GG allows for massive data collection, and it is expected that statistical analyses of the players’ hit rates can discriminate between explicit and implicit learning (EL and IL) impairment processes behind the players’ decision-making models. Assuming that IL system is the groundwork to build automaticity, it is plausible to suppose that GG is able to predict automaticity gait decline in people with PD. Then, the purpose of this study was to investigate the predictive power of GG in comparison with MoCA for gait automaticity impairments in PPD.

## 2 MATERIALS AND METHODS

### 2.1 Participants

A convenient sample of seventy-four PPD recruited from the Laboratory of Motor Learning participated in this study. Inclusion criteria involved were individuals with (1) idiopathic Parkinson’s disease as diagnosed by an experienced specialist in movement disorders, following the UK Brain Bank criteria [31], taking antiparkinsonian medications, (2) in 1-3 disease stage according to Hoehn and Yahr scale [28], (3) able to ambulate independently, (4) no signals of dementia (as determined by MoCA – cut-off 25) and/or major depression (as determined by Geriatric depression scale – cut-off 6). Subjects were excluded if they had clinically significant musculo-skeletal, cardio-vascular or respiratory disease, other neurological disease, or uncorrected visual/auditive disturbances.

### 2.2 Design and procedures

This study was approved by a Local Ethical Committee (CAAE 67388816.2.0000.0065) and conducted in accordance with the Helsinki Declaration. A written informed consent was signed for each participant before the study begun.

Based on a cross-sectional design, participants completed motor and cognitive evaluation in a single section. Evaluation order was randomized by sortition (Figure 1). Individual evaluation was conducted by a nurse and a physiotherapist specialized in movement disorders. All participants with PD were tested 40 to 120 minutes after their L-dopa dose (ON period).

**Figure 1:**
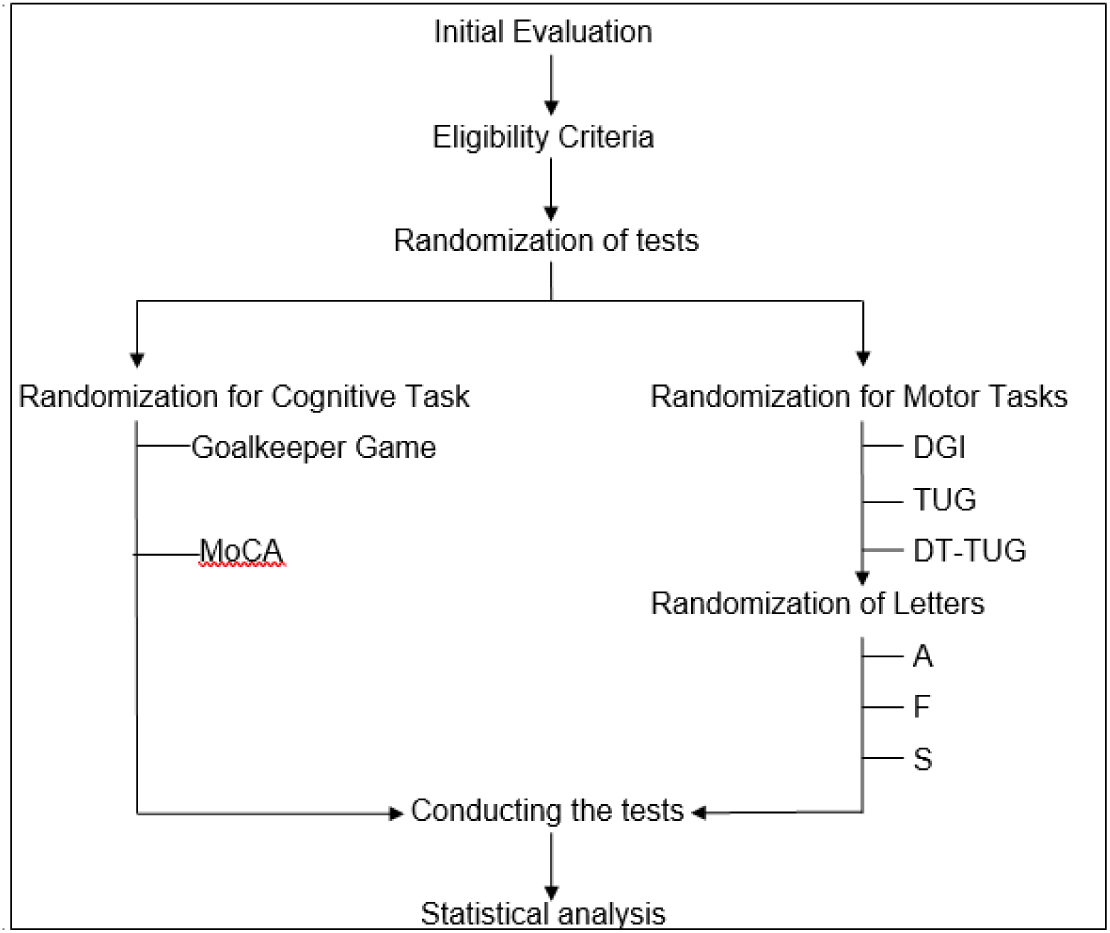
Study design flowchart.

### 2.3 Cognitive evaluation

Participants performed cognitive evaluation (GG and MoCA), seated comfortably in front of a desk where they could place elbows and forearms.

#### 2.3.1 GG evaluation

GG was presented in a 23” monitor (height = 29 cm, width = 51 cm) positioned 60 cm in front of the participants. After initial explanation about the game’s rules, participants are asked to assume the goalkeeper role during the penalty shootout, pressing the selected key among three possibilities (◂, ▾ or ▸) which controls the direction of the goalkeeper’s movement to defend the penalty (left, center or right).

The GG version used in this study is a simplified one with only deterministic penalty sequences. It has 3 phases: (1) MOTOR BASELINE PHASE, in which visual cues were offered to participants to guide the correct direction (5 trials); (2) IL PHASE, when no cues were offered and the sequences were provided by a deterministic model (20 trials); and (3) EL PHASE, in which participants were asked to memorize the correct sequence before playing the game (5 trials) (Figure 2).

**Figure 2:**
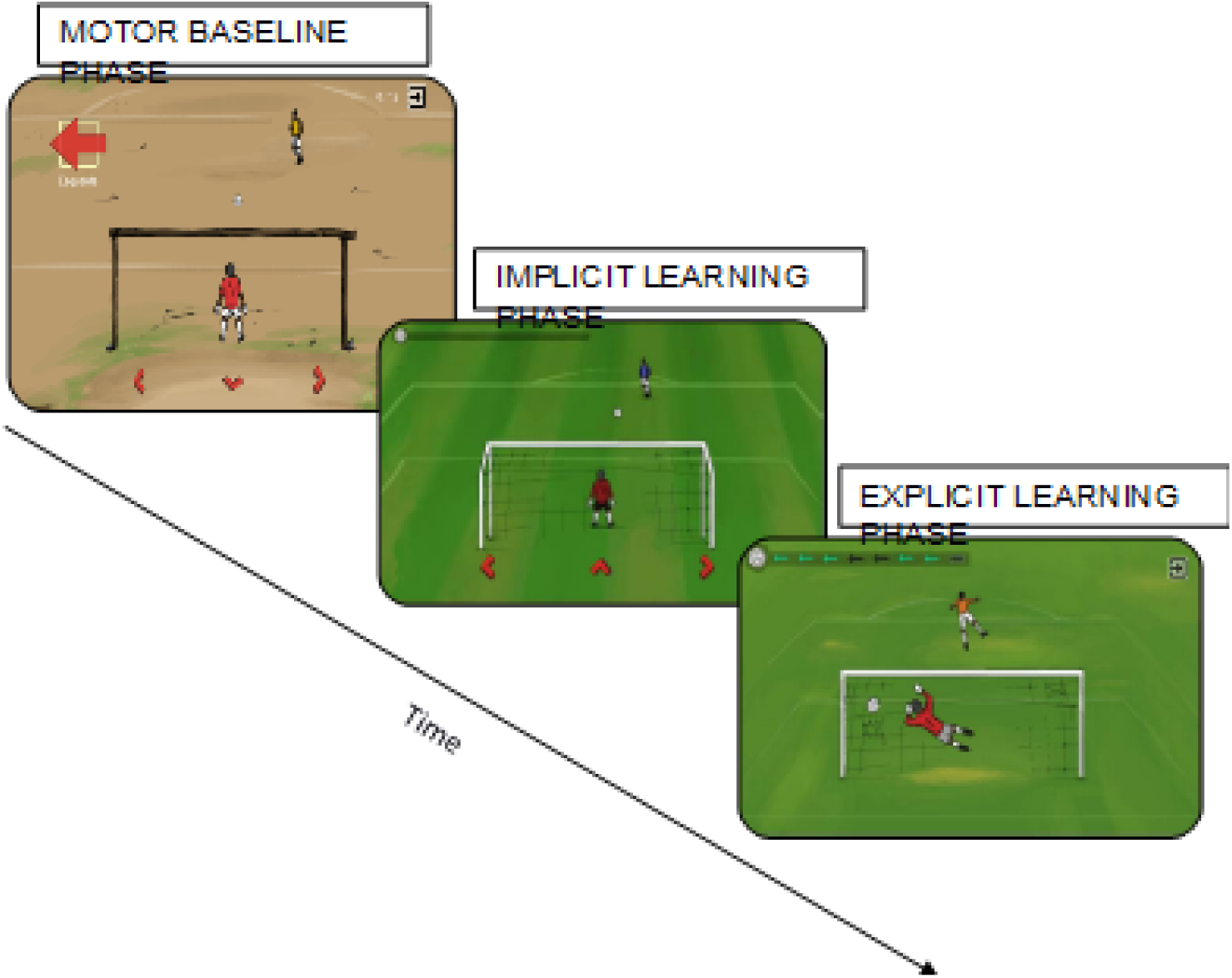
Goalkeeper Game where the participant was instructed to control the movements of a goalkeeper in time for a penalty, needing to guess the direction in which the opponent kicked the ball (left, right or center). At each level, the opponent adopted a new strategy based on implicit or explicit learning.

After every trial, participants received visual and auditory feedback indicating successful or unsuccessful attempt. All participants had to complete the three phases regardless of their performance.

#### 2.3.2 MoCA evaluation

The MoCA consists of 12 individual tasks that are scored and summed with a 6-item orientation screening and an educational correction (i.e., one point added for individuals with 12 years of education or less) to generate a total score representing global cognitive functioning. The total score is 30 points among seven cognitive domains, which reflect similar constructs as those assembled from a more comprehensive neuropsychological battery [60]. MoCA was administered with paper-and-pencil by a nurse with appropriate training. Participants received no feedback and completed the assessment regardless of their performance.

### 2.4 Motor evaluation

The motor evaluations were performed in a large room with adequate illumination and floor surface by a trained physiotherapist.

#### 2.4.1 Dynamic Gait Index (DGI)

This test assesses gait performance during 8 gait-related activities. These include quality of walking speed change, going around and over obstacles and stair walking as well as number of steps required for a pivot turn. Performance is scored from 0 to 24 indicating, respectively, lowest and highest functioning level [30]. DGI is highly recommended as a test to assess gait performance and has demonstrated good feasibility, test-retest and interrater reliability in PD. Furthermore, it is considerably useful as a supportive test for identifying the fall risk of people with PD [32]. After the initial explanation about the test, participants were asked to walk in habitual speed following examiner’s instructions.

#### 2.3.2 Dual-task Timed up to go test (DT-TUG)

TUG is a widely used test capable of providing valuable information on balance and mobility in PD [41, 49, 35]. This test measures the time in seconds that a patient takes to get up from a chair, walk 3m, make an 180*o* turn and go back to the original sitting position. Although walking is a large component of the test, its execution demands more than walking and it is unique in its serial integration of several mobility tasks. TUG has been considered an instrument with reliability and construct validity in PD [41, 4, 14] and is strongly recommended to assess gait performance and mobility in PD [5].

DT-TUG, i.e., TUG performed concomitantly with another task, has been used to increase TUG sensibility. DT-TUG is more sensitive than single-task TUG in identifying frail and prefrail individuals [36, 58], impairment in focused attention [10] and fallers [46] in elderly and PPD [56]. The concurrent cognitive task used in DT-TUG in this study consisted of speaking as many words as possible starting with a specific letter (F, S or A) presented at the beginning of test. After initial explanation, participants were asked to stand up from a chair, walk 3 m at normal speed, turn around, and come back to sit in the chair. Concomitantly, they had to complete a cognitive task (DT-TUG) or not (TUG). The tests order was randomized.

## 3 Statistical analysis

The GG variables cannot be compared directly to other clinical variables. For instance, for each stage and patient, the GG consists of a sequence of failures and successes in the patient’s predictions for that stage. Since each particular prediction carries little information, it is weakly correlated to the patient’s clinical variables.

In order to overcome this problem, we built a model which extracts each patient’s overall performance in the GG. This model is a generalization of logistic regression [17][p.119]. Specifically, for each time iteration, *t*, patient, *p*, and stage of GG, *s*, we define *X*_*t,p,s*_ as the indicator that *p* made the correct prediction at iteration *t* of stage *s* of the GG. The distribution of *X*_*t,p,s*_ is given by

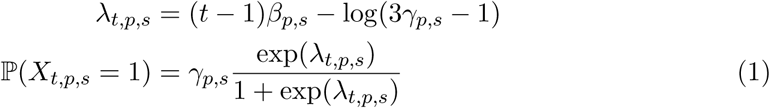

The model in eq. 1 admits an intuitive interpretation. First, it is chosen so that the probability that a patient makes a correct prediction at the first iteration of each stage is 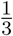. This is reasonable since, at that point, the patient has no information and there are 3 options. Also, the parameter *γ*_*p,s*_ represents the limit of patient *p*’s learning at stage *s*. That is, *γ*_*p,s*_ is the probability that *p* makes a correct prediction at *s* after playing that stage for a large number of iterations. Finally, *β*_*p,s*_ is the rate of learning of patient *p* at stage *s*. That is, *β*_*p,s*_ determines how many iterations *p* requires at stage *s* so that his probability of making a correct prediction is close to *γ*_*p,s*_.

Posterior estimates for *β*_*p,s*_ and *γ*_*p,s*_ were obtained using Stan [9]. As a result, three pairs of *β* and *γ* were attributed to each patient. By complementing these values with the average time spent per iteration in each stage, we obtained 9 variables that measure the patient’s performance in the GG.

We compared these variables and the MoCA score with respect to predictive power for clinical variables related to automaticity. Specifically, we built classifiers based on either the GG or the MoCA score to predict whether DGI and TUG scores where above or below the median. The classifiers were adjusted using elastic-net regularized logistic regression [18, 68]. The ROC curves for each pair of explanatory and response variable are presented below.

## 4 Results

Demographic and clinical features of participants are shown in Table 1.

**Table 1:** Demographic and clinical assessment data of the patients with Parkinson’s disease (n = 74). For continuous variables, the mean value is presented and also the standard deviation in parenthesis. For categorical variables, the mode is presented and also the mode’s proportion in parenthesis. GDS stands for the geriatric depression scale.

|  | HY 1 (n = 23) | HY 2 (n = 31) | HY 3 (n = 20) |
| --- | --- | --- | --- |
| Age (years) | 68.87 (9.39) | 63.55 (8.14) | 69.95 (8.67) |
| Gender (male) | M (14) | M (22) | M (17) |
| Education (years) | 11.30 (4.49) | 12.90 (5.31) | 15.40 (4.21) |
| UPDRS-III | 14.42 (7.35) | 19.71 (7.06) | 25.95 (11.34) |
| MoCA (total) | 24.78 (2.70) | 24.90 (3.43) | 24.25 (2.77) |
| *Visuospatial/executive* | 3.78 (1.13) | 3.58 (1.09) | 3.35 (1.46) |
| *Naming* | 2.65 (0.65) | 2.77 (0.43) | 2.85 (0.37) |
| *Attention* | 5.13 (0.97) | 5.03 (0.91) | 4.80 (0.95) |
| *Language* | 1.70 (0.88) | 1.94 (0.96) | 1.90 (0.91) |
| *Abstraction* | 1.87 (0.34) | 1.84 (0.45) | 1.80 (0.62) |
| *Memory (delayed recall)* | 3.43 (1.16) | 3.55 (1.06) | 3.15 (0.93) |
| *Orientation* | 5.57 (0.59) | 5.61 (0.80) | 5.60 (0.60) |
| GDS | 4.74 (3.00) | 4.06 (2.53) | 5.85 (3.23) |

The classifiers in the previous section show that GG variables are at least as good as MoCA for predicting DGI and TUG cost. This relation can be seen in Figure 3. On the one hand, GG variables are better at predicting DGI than MoCA, since the ROC curve for the former is higher than that for the latter: the estimated proportion of correct classifications for DGI with optimum cutoff is 65% using GG variables and 56% using MoCA. On the other hand, both GG variables and MoCA have low predictive power for TUG cost: their ROC curves are close to the identity line. When using optimum cutoffs, the estimated proportion of correct classifications for TUG cost using GG variables is 59% and using MoCA is 57%.

**Figure 3:**
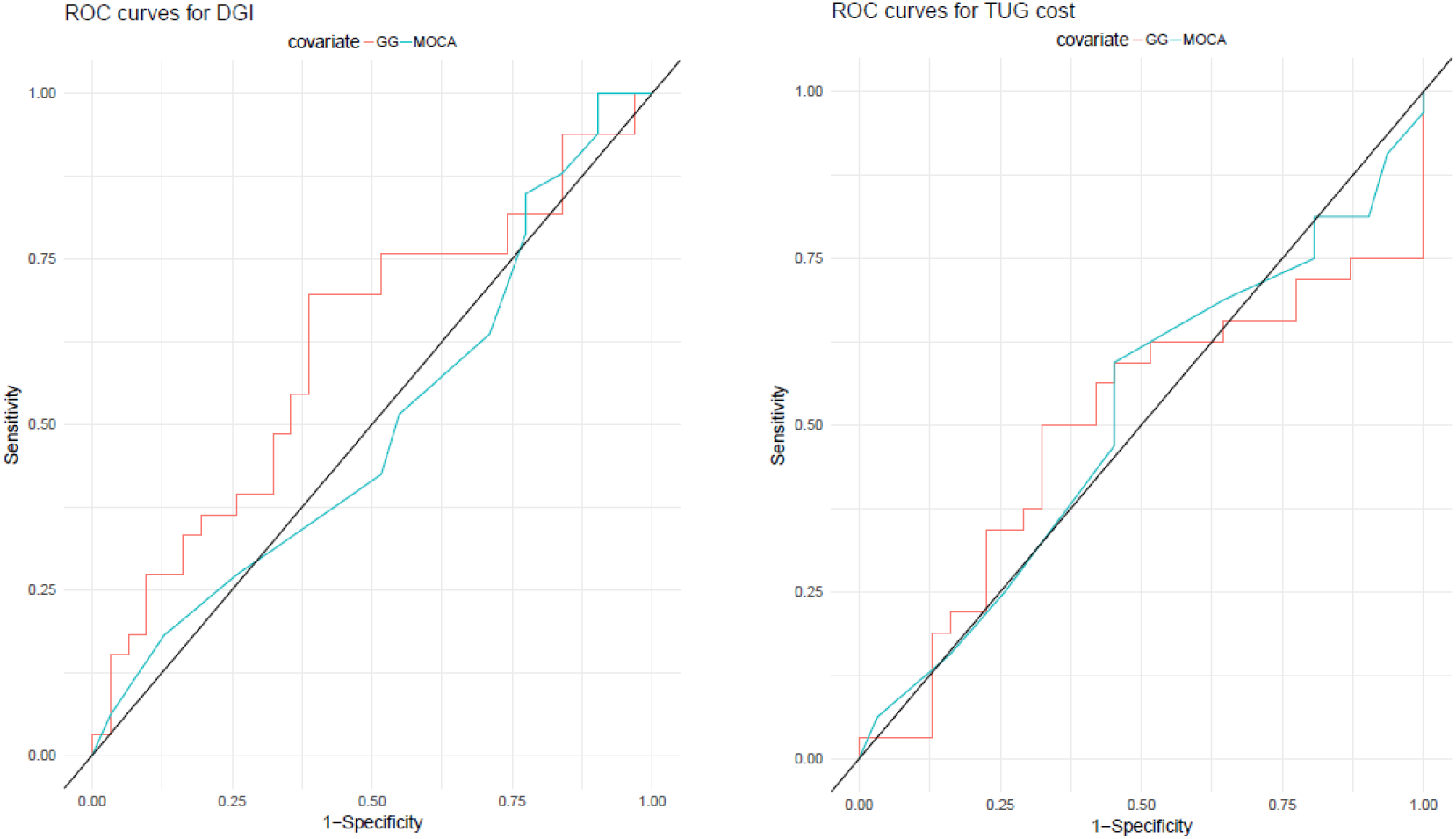
ROC (receiver operator characteristic) curves for predicting DGI (left) and DT-TUG cost (right) using the classifiers adjusted with either GG variables or MoCA score.

## 5 Discussion

The main finding of this study was that GG has higher power to predict decline in gait performance under complex situations and DT conditions than MoCA. MoCa is a recommended and popular test in PD36, however, as shown elsewhere, it fails to detect gait decline under DT [20]. In face of this, the main purpose of this study was to test whether GG can be used under DT to predict disruption in gait automaticity in early stages of PD evolution. GG allows the use of several strategy trees, deterministic and non-deterministic, which can be tailored to specific experimental protocols.

The GG version adopted in this work was chosen with the purpose of achieving a friendly and free tool able to identify IL deficit regardless of motor control and EL deficits. Thus, the game was designed with 3 phases. The first of them obtains the motor performance baseline, the second evaluates the ability to identify the direction sequence by IL only, and the third evaluates the ability to identify the direction sequence by EL. The motor deficits associated to, e.g., rest tremor or movement slowness, could be isolated as also the possible EL or attentional deficits associated to other cognitive alterations.

IL is a knowledge acquiring process that occurs without conscious awareness of learn ing, whereas EL involves the use of overt strategies. Seminal studies showed that people with amnesia exhibited normal learning of the motor task but had severely impaired EL. In contrast, PPD failed to learn probabilistic classification tasks, despite having intact EL. This double dissociation showed that the limbic-diencephalic regions, damaged in amnesia, and the neostriatum damaged in PD, support separate and parallel learning systems [52, 34]. Then, the well-functioning of neostriatum circuits is essential for the gradual, incremental learning of associations that is characteristic of habit learning. In other words, the well-functioning of neostriatum allows the construction of automaticity, i.e., the control of a learned task with minimal attentional demand [64, 48, 24].

Other more recent studies in healthy individuals [16] and PPD have confirmed that neostriatum circuits are associated to implicit sequence learning [62, 54, 23, 26] and probabilistic implicit sequences learning [61, 63, 21].

Therefore, as expected, a tool developed to be sensitive to implicit sequence learning, isolating the influence of motor and EL components, had to be more efficient than a general cognitive test as MoCA to predict the decline in automaticity associate to PD, a disease that has as main pathogenic mechanism dopamine depletion in basal ganglia circuits. Besides, although the comparison of the participants’ engagement in doing the GG and MoCA tests was not a purpose of the present study, apparently GG was able to arise more their motivation, imposing a lower level of stress than a paper-and-pencil test as MoCa.

On the other hand, the two gait tests used to assess gait automaticity imposed challenging gait conditions for participants where the cognitive resources are more demanded than only walking. Previous studies showed a higher activation in prefrontal cortex during obstacle negotiation in comparison to no obstacle in the gait in young and older adults [39] and PwPD [37]. Even in young healthy individuals the performance level in TUG declined under DT conditions [57]. Taken together, this evidence corroborate that more complex gait conditions demand more cognitive resources and, therefore, are more sensitive to assess gait automaticity than only walking. Besides, the movement sequences tested in TUG and obstacle negotiation tested in DGI are frequently performed in daily life and the performance in these tests is associated with fall risk.

Therefore, it is plausible to hypothesize that GG will be able to predict the level of impairment and fall risk in daily living activity in PPD. To answer this, novel studies should be conducted.

Currently, it has been proposed that early PD could be divided into 3 stages: preclinical in which the neurodegenerative process is started, without evident symptoms or signs of the disease; prodromal, in which symptoms and signs of this disease are present but are insufficient to define a full clinical picture; and clinical, in which the diagnosis is achieved, based on the presence of classical motor signs. Prodromal disease refers to the stage wherein early symptoms or signs of PD neurodegeneration are present, but a clinical diagnosis is not yet possible. Finding new criteria to identify prodromal PD represents a promising challenge for research in PD [38]. In this direction, further studies should be conducted to investigate the ability of GG to identify early decline in gait automaticity in prodromal PD stages.

Finally, it is fundamental that we point some limitations of the present study for result generalization. These are: the small sample size, the absence of a control group and the absence of another test considered as “gold standard” for IL in PD, allowing comparison of results. The deterministic version of game used in our study was able to assess aspects of cognitive tasks performances, making it a tool comparable to MoCA tests for PD. With the development of more strategy trees, including some non-deterministic, and new protocols, it is expected the GG will become a powerful tool in PD screening.

## 6 Acknowledgments

This paper stems from research activity conducted as part of the Research, Innovation and Dissemination Center for Neuromathematics (NeuroMat), funded by the São Paulo Research Foundation-FAPESP (grant 2013/07699-0). MD and YLU are supported by CAPES scholarships. ACR is also recipient of a Brazilian National Council for Scientific and Technological Development-CNPq research productivity grant (# 306251/2014-0). We thank Antonio Galves for stimulating discussions.

